# Multimodal Neuroimaging-based Prediction of Adult Outcomes in Childhood-onset ADHD using Ensemble Learning Techniques

**DOI:** 10.1101/785766

**Authors:** Yuyang Luo, Tara L. Alvarez, Jeffrey M. Halperin, Xiaobo Li

**Author notes:** **Correspondence**: Xiaobo Li, PhD, Associate Professor of Biomedical Engineering, New Jersey Institute of Technology, 323 Martin Luther King Blvd, Fenster Hall 613, Newark, NJ 07102, **Email:**.

## Abstract

Attention deficit/hyperactivity disorder (ADHD) is a highly prevalent and heterogeneous neurodevelopmental disorder, currently relaying on subjective symptom observations for diagnosis. Machine learning classifiers have been utilized to assist the development of neuroimaging-based biomarkers for objective diagnosis of ADHD. However, the existing basic model-based studies in ADHD reported suboptimal classification performances and inconclusive results, mainly due to the limited flexibility for each type of basic classifiers to appropriately handle multi-dimensional source features with various properties. In this study, we proposed to apply ensemble learning techniques (ELTs) in multimodal neuroimaging data collected from 72 young adults, including 36 probands (18 remitters and 18 persisters of childhood ADHD) and 36 group-matched controls. All the currently available optimization strategies for ELTs (i.e., voting, bagging, boosting and stacking techniques) were tested in a pool of semi-final classification results generated by seven basic classifiers. The high-dimensional neuroimaging features for classification included regional cortical gray matter (GM) thickness and surface area, GM volume of subcortical structures, volume and fractional anisotropy of major white matter fiber tracts, pair-wise regional connectivity and global/nodal topological properties of the functional brain network for cue-evoked attention process. As a result, the bagging-based ELT with the base model of support vector machine achieved the best results, with the most significant improvement of the area under the receiver of operating characteristic curve (0.89 for ADHD vs. controls, and 0.9 for ADHD persisters vs. remitters). We found that features of nodal efficiency in right inferior frontal gyrus, right middle frontal (MFG)-inferior parietal (IPL) functional connectivity, and right amygdala volume significantly contributed to accurate discrimination between ADHD probands and controls; higher nodal efficiency of right MFG greatly contributed to inattentive and hyperactive/impulsive symptom remission, while higher right MFG-IPL functional connectivity strongly linked to symptom persistence in adults with childhood ADHD. Our study also suggested that considering their solidly improved robustness than the commonly implemented basic classifiers, ELTs may have the potential to identify more reliable neurobiological markers for severe brain disorders.

## Introduction

Attention deficit/hyperactivity disorder (ADHD) is a heterogeneous neurodevelopmental disorder. Over the past two decades, the estimated prevalence of diagnosed ADHD among the U.S. children and adolescents has significantly increased from 6.1% to 10.2% (Xu et al., 2018). The current ADHD diagnostic standards are fully clinical symptom-based, and relay on information collected from multiple sources through subjective observations, which often cause biases and inconsistencies of the diagnoses.

Multimodal neuroimaging techniques have been widely implemented to investigate the neural mechanisms of ADHD. A number of structural MRI and Diffusion Tensor Imaging (DTI) studies have suggested that gray- and/or white-matter (GM/WM) structural underdevelopment in frontal lobe, thalamus, and striatum significantly contribute to the emergence of ADHD during childhood (Ellison-Wright et al., 2008; Xia et al., 2012). Meanwhile, functional aberrances in the fronto-thalamo/fronto-striatal circuitries have also been frequently reported to link with symptom onset in children with ADHD. For instance, altered task-driven or spontaneous neural activities in prefrontal cortex, thalamus, and striatum, and their functional connectivities, have been found to significantly associate with increased inattentive and/or impulsive symptoms in children with ADHD (Bush et al., 2005; Cubillo et al., 2012; Durston, 2003; Li et al., 2012; Rubia et al., 1999; Yang et al., 2011). Simultaneously, increasing neuroimaging studies have found that optimal structural/functional development in fronto-subcortical pathways may contribute to symptom reduction and remission of ADHD in adulthood. For instance, a longitudinal study found that persistently decreased GM thickness in dorsolateral prefrontal, middle frontal, and inferior parietal regions, and reduced WM fractional anisotropy (FA) in left uncinate and inferior frontal-occipital fasciculi were associated with a greater number of ADHD symptoms persisting into adulthood (Shaw et al., 2013; Shaw et al., 2015). Proal et al. reported that adults with persistent ADHD had thinner cortical thickness relative to the remitted ADHD in prefrontal region (Proal et al., 2011). In addition, greater prefronto-thalamo functional connectivity during cue-evoked attention process (Clerkin et al., 2013), and greater within-frontal functional connectivity during resting-state (Francx et al., 2015), have been observed in adult ADHD remitters relative to the persisters.

However, findings of the entire body of existing neuroimaging studies are widely inconsistent, partially due to the sample biases, differences of the implemented imaging and analytic techniques, and the limitations of the traditional parametrical models for group comparisons. Indeed, traditional statistical methods (e.g. t-tests, analysis of variance (ANOVA), correlation, etc.) estimate group differences only within a voxel or region of interest (ROI) at a time without having the capacity to explore how ROIs interact in linear and/or non-linear ways, as they quickly become overburdened when attempting to combine predictors and their interactions from high dimensional imaging data sets (Sun et al., 2009).

Compared to traditional parametrical models, multivariate machine learning techniques are able to leverage high dimensional information simultaneously to understand how variables jointly distinguish between groups (Greenstein et al., 2012). In literature, support vector machine (SVM) is the most frequently applied machine learning classifier in neuroimaging data from children with ADHD, which has been aided by recursive feature elimination (RFE), temporal averaging, principle component analysis (PCA), fast Fourier transform (FFT), independent component analysis (ICA), 10-fold cross-validation (CV), hold-out, and leave-one-out cross-validation (LOOCV) techniques, to distinguish children with ADHD from normal controls (Brown et al., 2012; Chang et al., 2012; Cheng et al., 2012; Colby et al., 2012; Du et al., 2016; Fair et al., 2012; Iannaccone et al., 2015; Johnston et al., 2014; Sen et al., 2018; Yasumura et al., 2017). The commonly reported most important features (according to importance score) that contribute to successful group discrimination included functional connectivity of bilateral thalamus, functional connectivity, surface area, cortical curvature and/or voxel intensity in frontal lobe, cingulate gyrus, temporal lobe, etc. (Brown et al., 2012; Colby et al., 2012; Iannaccone et al., 2015). SVM has also been applied to structural MRI and DTI data collected from adults with ADHD and controls, which reported between-group differences in widespread GM and WM regions in cortices, thalamus, and cerebellum (Chaim-Avancini et al., 2017). Meanwhile, neural network-based techniques, including deep belief network, fully connected cascade artificial neural network, convolutional neural network, extreme learning machine, and hierarchical extreme learning machine, have also been utilized to structural MRI and resting-state functional MRI (fMRI) data in children with ADHD and controls (Deshpande et al., 2015; Kuang and He, 2014; Peng et al., 2013; Qureshi et al., 2016; Qureshi et al., 2017; Zou et al., 2017). The most important group discrimination predictors identified by these neural network studies included functional connectivities within cerebellum, functional connectivity, surface area, cortical thickness and/or folding indices of frontal lobe, temporal lobe, occipital lobe and insula (Deshpande et al., 2015; Peng et al., 2013; Qureshi et al., 2017). In addition, principle component-based Fisher discriminative analysis (PC-FDA) (Zhu et al., 2008), Gaussian process classifiers (GPC) (Hart et al., 2014; Lim et al., 2013), and multiple kernel learning (Dai et al., 2012; Ghiassian et al., 2016) have also been used in functional and structural MRI data to discriminate children with ADHD from controls. More details of existing machine learning studies in ADHD are provided in **Supplementary Table 1**. These existing studies have either utilized features representing regional/voxel brain properties collected from only single imaging modality, or the combination of two modalities (mostly structural MRI and resting-state fMRI) (Brown et al., 2012; Fair et al., 2012; Hart et al., 2014; Iannaccone et al., 2015; Johnston et al., 2014), or reported poor accuracy (Dai et al., 2012; Sen et al., 2018; Zou et al., 2017). Some studies did not conduct the very necessary step of estimating the most important features that contribute to accurate classifications (Chang et al., 2012; Dai et al., 2012; Kuang and He, 2014; Qureshi et al., 2016; Sen et al., 2018; Tenev et al., 2014; Zou et al., 2017). In this field, systems-level functional and structural features, such as global and regional topological properties from functional brain networks during cognitive processes and WM tract properties have not been considered. In addition, relations between the suggested predictors from imaging features and clinical/behavioral symptoms in samples of ADHD patients, which can provide important clinical context, have not been studied.

Ensemble learning techniques (ELTs), which propose to integrate results from multiple basic classifiers by using voting (Lam and Suen, 1997; Ruta and Gabrys, 2005), bagging (Breiman, 1996), stacking (Wolpert, 1992), or boosting (Johnston et al., 2014; Schapire, 1990a; Yoav Freund and Schapire, 1997) strategies, have been recently developed in the big data science field, to deal with complicated feature variations, biases, and optimize prediction performances (Deng and Platt, 2014; Wang et al., 2011). From literature, ELTs have only been applied in three recent studies to discriminate patients with ADHD from controls (Eloyan et al., 2012; Tenev et al., 2014; Zhang-James et al., 2019). Eloyan and colleagues applied a voting-based ELT, along with hold-out technique for CV, in structural MRI and resting-state fMRI data from children with ADHD and controls, and reported an important group discrimination predictor of dorsomedial-dorsolateral functional connectivity in the motor network (Eloyan et al., 2012). Voting-based ELT has also been applied in electroencephalogram (EEG) data collected from adults with ADHD and controls, without reporting the most important discrimination predicators (Tenev et al., 2014). Very recently, a preprint from Zhang-James et al. conducted a study that applied ELTs in structural MRI data from patients with ADHD (both adults and children) and controls, and suggested that GM volume of bilateral caudate and thalamus and orbitofrontal surface area significantly contribute to successful group discrimination (Zhang-James et al., 2019). However, the preprint was lacking of clarifications about optimization strategies and reported low accuracy of discriminations (<0.65).

In the current study, we proposed to apply ELTs in structural MRI, DTI, and task-based fMRI data collected from a sample of adults with childhood ADHD and controls, who have been clinically followed up since childhood. All the currently available optimization strategies (i.e., voting, bagging, boosting and stacking techniques) were tested in a pool of semi-final classification results generated by seven basic classifiers (including K-Nearest Neighbors (KNN), SVM, logistic regression (LR), Naïve Bayes (NB), linear discriminant analysis (LDA), random forest (RF), and multilayer perceptron (MLP)). A nested CV including an inner LOOCV and an outer 5-fold CV were applied with grid search to tune the hyperparameters and minimize the overfitting. The high-dimensional neuroimaging features for classification included regional cortical GM thickness and surface area, GM volume of subcortical structures estimated from structural MRI data, volume and FA of major WM fiber tracts derived from DTI data, the pair-wise regional connectivity and global/nodal topological properties (i.e., global-, local-, and nodal-efficiency, etc.) of the cue-evoked attention processing network computed from task-based fMRI data. Based on findings from existing studies from our group and other teams, we hypothesized that structural and functional alterations in frontal, parietal, and subcortical areas and their interactions would significantly contribute to accurate discrimination of ADHD probands (adults diagnosed with ADHD in their childhood) from controls; while abnormal fronto-parietal hyper-communications in right hemisphere would play an important role in inattentive and hyperactive/impulsive symptom persistence in adults with childhood ADHD. In addition, we hypothesized that the classification performance parameters (accuracy, area under the curve (AUC) of the receiver operating characteristics (ROC), etc.) of ELTs-based procedures would be greatly improved compared to those of basic model-based procedures.

## Materials and methods

### Participants

A total of 72 young adults (age mean=24.4 years, standard deviation=2.1 years), who provided complete and good quality data from multimodal neuroimaging and clinical assessments, were involved in this study. There were 36 ADHD probands diagnosed with ADHD combined-type (ADHD-C) in their childhood and 36 group-matched comparison subjects with no history of ADHD. Among the 36 ADHD probands, 18 were classified as ADHD remitters, who were endorsed no more than 3 inattentive or 3 hyperactive/impulsive symptoms in adulthood and had no more than 5 symptoms in total. The rest of 18 probands were classified as ADHD persisters, endorsed at least five inattentive and/or hyperactive/impulsive symptoms in their adulthood and had at least 3 symptoms in each domain.

All the 72 subjects involved in this study had been clinically follow-up since their 7-10 years of ages. Childhood diagnoses were based on teacher ratings using the IOWA Conners’ Teachers Rating Scale (Loney and Milich, 1982) and parent interview using the Diagnostic Interview Schedule for Children version 2 (Shaffer et al., 1989). The exclusion criteria in childhood were chronic medical illness; neurological disorder; diagnosis of schizophrenia, autism spectrum disorder, or chronic tic disorder; Full Scale IQ < 70; and not speaking English. Adult psychiatric status was assessed using the Structured Clinical Interview for DSM-IV Axis I Disorders (First et al., 2002), supplemented by a semi-structured interview for ADHD that was adapted from the Schedule for Affective Disorders and Schizophrenia for School-Age Children (Kaufman et al., 1997) and the Conners’ Adult ADHD Diagnostic Interview for DSM-IV (Epstein et al., 2006). The raw scores of inattentive and hyperactive/impulsive symptoms from the Conner’s Adult Self-Rating Scale were normalized into T scores based on DSM-IV standard, and were used as dimensional measures for inattentive and hyperactive/impulsive behaviors. Exclusion criteria in adulthood were psychotropic medication that could not be discontinued and conditions that would preclude MRI (e.g., metal in body, pregnancy, too obese to fit in scanner). Clinical and demographic information are listed in Table 1.

**Table 1:** Demographic and clinical characteristics in groups of controls and ADHD probands (and further in the sub-groups of remitters and persisters of the ADHD probands).

|  | **Controls**<br>**(N = 36)** | **ADHD**<br>**(N = 36)** |  | **Remitted**<br>**(N=18)** | **Persistent**<br>**(N=18)** |  |
| --- | --- | --- | --- | --- | --- | --- |
|  | **Mean (SD)** | **Mean (SD)** | **p** | **Mean (SD)** | **Mean (SD)** | **p** |
| **Age** | 24.3 (2.3) | 24.66 (2.0) | 0.48 | 24.79 (2.2) | 24.52 (2.0) | 0.7 |
| **Full-scale IQ** | 103.83(15.4) | 97.96 (14.1) | 0.1 | 99.22 (14.9) | 96.71 (13.6) | 0.6 |
| **Conners' Adult ADHD**<br>**Rating Scale (T score)** |  |  |  |  |  |  |
| Inattentive | 45.75 (8.8) | 56.5 (13.2) | <0.001 | 49.83 (10.9) | 63.17 (12.0) | 0.001 |
| Hyperactive/impulsive | 42.97 (6.2) | 53.64 (12.9) | <0.001 | 46.17 (9.0) | 61.11 (12.0) | <0.001 |
| ADHD Total | 43.89 (8.2) | 56.5 (14.7) | <0.001 | 42.61 (7.5) | 54.33 (8.8) | <0.001 |
| **ADHD semistructured**<br>**interview (number of**<br>**symptoms)** | 0.79 (1.6) | 6.17 (5.2) | <0.001 | 2.64 (2.0) | 10.24 (3.6) | <0.01 |
|  | **N (%)** | **N (%)** | **p** | **N (%)** | **N (%)** | **p** |
| **Male** | 31 (86.1) | 30 (83.3) | 0.74 | 16 (88.9) | 14 (77.8) | 0.37 |
| **Right-handed** | 32 (88.9) | 32 (88.9) | 1 | 15 (83.3) | 16 (88.9) | 0.63 |
| **Race** |  |  | 0.17 |  |  | 0.59 |
| Caucasian | 15 (41.7) | 21 (58.3) |  | 9 (50.0) | 12 (66.7) |  |
| African American | 13 (36.1) | 7 (19.4) |  | 4 (22.2) | 3 (16.7) |  |
| More than one race | 6 (16.7) | 8 (22.2) |  | 5 (27.8) | 3 (16.7) |  |
| Asian | 2 (5.6) | 0 (0.0) |  | 0 (0.0) | 0 (0.0) |  |
| **Ethnicity** |  |  | 0.09 |  |  | 0.74 |
| Hispanic/Latino | 10 (27.8) | 17 (47.2) |  | 8 (44.4) | 9 (50.0) |  |
|  | **Mean (SD)** | **Mean (SD)** | **p** | **Mean (SD)** | **Mean (SD)** | **p** |
| **Task performance**<br>**measures** |  |  |  |  |  |  |
| Reaction time average | 395.8 (53.1) | 422.8 (74.3) | 0.08 | 431.1 (67.0) | 439.1 (107.8) | 0.79 |
| Reaction time std | 129.6 (24.8) | 137.2 (29.9) | 0.25 | 136.2 (27.6) | 138.2 (32.8) | 0.84 |
| Anticipation error | 1.86 (2.1) | 1.74 (1.6) | 0.78 | 1.69 (1.6) | 1.78 (1.7) | 0.88 |
| Commission error | 0.33 (0.8) | 0.85 (1.4) | 0.07 | 0.75 (1.6) | 0.94 (1.3) | 0.7 |
| Omission error | 4.97 (5.8) | 8 (10.8) | 0.15 | 4.38 (4.0) | 11.22 (13.8) | 0.06 |

The study received Institutional Review Board approval at the participating institutions. Participants provided signed informed consent and were reimbursed for their time and travel expenses.

### Multimodal Neuroimaging Data Acquisition Protocol

Multimodal neuroimaging data of each participant were collected using the same 3.0T Siemens Allegra (Siemens, Erlangen, Germany) whole body MRI scanner. High resolution 3-dimensional T1-weighted structural MRI data was acquired using magnetization prepared rapid gradient echo pulse sequence with TR=2050 ms, TE=4.38 ms, inversion time (TI)=1.1 s, flip angle=8°, field of view (FOV)=256 mm×256 mm×256 mm, voxel size=0.94 mm×0.94 mm×1 mm. DTI data was collected using an echo planar imaging (EPI) pulse sequence with a b-value=1250 s/mm^2^ along 12 independent non-collinear orientations, as well as one reference volume without diffusion-weighting b=0 s/mm^2^ (TR=5200 ms, TE=80 ms, flip angle=90°, FOV=128 mm×128 mm, voxel size=1.875 mm×1.875 mm×4 mm, matrix=128×96, number of slices=63). FMRI data were acquired using a gradient-echo EPI sequence with TR=2500 ms, TE=27 ms, flip angle=82°, matrix=64×64, slice thickness=4 mm, 40 slices, in-plane resolution=3.75 mm^2^. Images were acquired with slices positioned parallel to the anterior commissure-posterior commissure line. The total duration of fMRI data acquisition was 20 minutes, which contained 4 runs of a cued attention task (CAT) with stimuli counter-balanced.

### The CAT for fMRI

The CAT was developed and described in details in (Clerkin et al., 2013; Clerkin et al., 2009; Luo et al., 2018). Briefly, it began and ended with a 30-second fixation period, respectively. The task contained a series of 120 letters, including 24 targets (“X”), 12 cues (“A”), and 84 non-cue letters (“B” through “H”). For the 24 targets, half of them preceded by a cue, and the other half preceded by a non-cue letter. Participants were told that a cue letter was always followed by a target letter, but not all targets were preceded by a cue letter. The stimuli of letters were presented individually for 200 ms with a pseudorandom inter-stimulus interval which ranged from 1550 to 2050 ms (mean=1800 ms/run). Participants were instructed to respond to each target as rapidly as possible using their right index finger. Before entering into the scanner, very detailed instructions and practical trials of the task were provided to each participant to ensure satisfied performance.

### Multimodal Imaging Data Processing for Feature Extractions

The T1-weighted data were reconstructed into a 3-dimensional cortical model for thickness and area estimations using FreeSurfer v.5.3.0 (https://surfer.nmr.mgh.harvard.edu). Each volume was first registered to the Talairach atlas to compute the transformation matrix using an affine registration method. Then intensity variations caused by magnetic field inhomogeneities were corrected and non-brain tissue was removed. In addition, a cutting plane was used to separate the left and right hemispheres and to remove the cerebellum and brainstem. Two mess surfaces (mess of grids created using surface tessellation technique) were generated between GM and WM (white matter surface), as well as between GM and cerebrospinal fluid (pial surface). The distance between the two closest vertices of the WM and pial surfaces presented the cortical thickness at that specific location. Regional cortical thickness and area in 68 bilateral cortical regions were estimated based on the Desikan atlas. Based on a manually labelled model constructed according to prior knowledge of spatial relationships acquired with a training data set, volume of each subcortical structure was then calculated for each subject. At the end, a total of 202 structural MRI features, including regional cortical GM thickness, surface area, and GM volume of subcortical structures were extracted from each subject.

For each subject, the DTI data was corrected for eddy current-induced distortions due to the changing gradient field directions. Meanwhile, head motion was corrected with non-diffusion-weighted reference image (b0 image) using an affine, 12 degrees of freedom registration. Then the FA value and principle diffusion direction at each brain voxel were calculated. To conduct WM probabilistic tractography between each pair of 18 ROIs (including thalamus, putamen and caudate nuclei from striatum, hippocampus, and frontal, parietal, occipital, temporal, and insular cortices in both hemispheres), the FSL/BEDPOSTX tool was used to generate the probability distribution of diffusion parameters within each voxel, including modeling for diffusion of crossing fibers along two directions. These 18 ROIs were creased based on the Harvard-Oxford Cortical Atlases and the Julich Histological Atlas from the MNI standard space, and mapped to the DTI data. We used the multi-fibre probabilistic connectivity-based method to determine the number of pathways between the seed and each of the target clusters. The default setting of parameters for the Markov Chain Monte Carlo estimation of the probabilistic tractrography was utilized: 5000 individual pathways were drawn on the principle fiber direction of each voxel within the seed ROI using the probability distributions; curvature threshold of 80° to exclude implausible pathways; a maximum number of 2000 travel steps of each sample pathway and a 0.2 mm step length. Therefore, a total of 120 DTI properties, including the volume and FA of cortico-cortical and subcortico-cortical WM fiber tracts were extracted for each subject.

The fMRI data from each participant was preprocessed using Statistical Parametric Mapping version 8 (SPM8, Wellcome Trust Centre for Neuroimaging, London, United Kingdom; http://www.fil.ion.ucl.ac.uk/spm/) implemented on a MATLAB platform. The preprocessing procedures included slice timing correction, realignment, coregistration, segmentation, normalization, and spatial smoothing. The first-level analyses were conducted using general linear model (GLM) to generate the activation map responding to the cues. The group average activation maps for ADHD probands and normal controls were generated, respectively. A total of 52 cortical and subcortical seed regions, which was parceled according to the structural and functional connectivity-based Brainnetome atlas, were determined based on the results of the combination of the functional activation maps of the groups of ADHD probands and controls (Fan et al., 2016). To construct the cue-evoked attention processing network, the single-trial beta value series from the 48 cue-related events in the four runs were extracted. Among all the voxels in each of the 52 node ROIs, the average beta value series was calculated and used to create a 52 × 52 pair-wise Pearson correlation matrix. Then the graph theoretic techniques (GTTs) were carried out. More details of the fMRI data processing can be found in (Luo et al., 2018). A total of 200 fMRI features, including the global- and local-efficiency of the entire network, the nodal efficiency, degree, and betweenness-centrality measures of the 52 nodes, as well as their pair-wise functional connectivities, were generated for each subject.

### Modeling of Ensemble Learning Architecture

Modeling of the ELTs for classifications between ADHD probands and controls, as well as between ADHD persisters and remitters respectively, was described in Figure 1. Specifically, **Part A** of Figure 1 presented feature selection and preparation flow. In order to decrease the risk of overfitting, two-sample t-tests were applied and a total of 20 neuroimaging features that showed the most significant between group differences were first selected from the 522 multimodal neuroimaging features derived from structural MRI, DTI, and fMRI data. Then normalized value of each of the 20 selected features were carried out by doing a z-score normalization in the feature-specific space. The normalized 20 top-ranking neuroimaging features were then entered to the training and validation procedures (**Part B** of Figure 1), which consisted of a nested CV (there were two CV loops, including an outer 5-fold CV loop to split the data into training set and validation set, and an inner loop to tune the hyperparameters for 7 basic models and 4 ELTs-based models using grid search in combination with LOOCV). More specifically, the 20 neuroimaging features were first split into a total of 5 stratified folds that each fold consisted of balanced 20% of the entire data. The five-fold CV was performed by using these 5 stratified folds, where each trial dedicated four folds for training data and the remaining one for validation. Then for each iteration in 5-fold CV, the corresponding training set was sent into the LOOCV processing. In each iteration, one subject was extracted from the training set to act as a validation data, and the remaining subjects were trained to construct the models. According to the classification performance of the validation data, the hyperparameters for each model were tuned and the optimal hyperparameters setup were selected using grid search. More details of the hyperparameters were described in Table 2. In the current study, we utilized the LOOCV to tune the hyperparameters of 7 basic models, including KNN, SVM, LR, NB, LDA, RF, and MLP. Based on the hyperparameters of basic models, we applied LOOCV to tune the hyperparameters of 4 ELTs-based models, including max Voting, Bagging, AdaBoost, Stacking. As shown in **Part C** of Figure 1, during iterations of 5-fold CV outer loop, the performance of each basic and ELTs-based models with the optimal hyperparameters derived from LOOCV inner loop iterations was evaluated. The group average of classification performance of each classifier derived from each iteration of 5-fold CV was generated. The 7 basic and 4 ELTs-based models according to the group average value of AUC of the ROC from iterations of 5-fold CV outer loop. The basic and ELTs-based models with the highest average AUC were selected as optimal classifiers. Based on the types of ELTs-based models we have evaluated and selected, the importance score corresponding each feature was then calculated using the ELTs-based model and the corresponding basic models.

**Table 2:** The hyperparameters of 7 basic models and 4 ELTs-based models.

| **Classifiers** | **Hyperparameters** |
| --- | --- |
| KNN | n_neighbors: [1, 3, 5, 7, 9]; algorithm: ['auto', 'ball_tree', 'kd_tree', 'brute']; p: [1, 2, 3] |
| SVM | C: [0.001, 0.01, 0.1, 1, 10, 100, 1000]; gamma: ['auto', 'scale']; kernel: ['linear', 'rbf', 'poly', 'sigmoid] |
| LR | solver: ['newton-cg', 'lbfgs', 'sag', 'saga']; multi_class: ['ovr', 'multinomial', 'auto'] |
| RF | n_estimators: list(range(3, 60, 5)); criterion: ['gini', 'entropy']; min_samples_leaf: [3, 5, 10];<br>max_depth: [3, 4, 5, 6]; min_samples_split: [3, 5, 10]; bootstrap: [True, False] |
| LDA | solver: ['svd', 'lsqr', 'eigen'] |
| MLP | activation: ['identity', 'logistic', 'tanh', 'relu']; solver: ['lbfgs', 'sgd', 'adam'];<br>hidden_layer_sizes: np.arange(1, 72, 10); max_iter: [4000] |
| ELT-Voting | estimators; voting: ['hard', 'soft'] |
| ELT-Bagging | base_estimator; n_estimators: list(range(10, 150, 10)); max_samples=[0.2, 0.3, 0.4, 0.5];<br>max_features=[0.5, 0.6, 0.7, 0.8, 0.9, 1.0] |
| ELT-Boosting | base_estimator; n_estimators: list(range(10, 150, 10)); learning_rate: list(range(0.01, 1, 0.01)) |
| ELT-Stacking | classifiers; meta_classifiers |

ELTs: ensemble learning techniques; KNN: k-nearest neighbors; SVM: support vector machine; LR: logistic regression; RF: random forest; LDA: linear discriminant analysis; MLP: multilayer perceptron.

**Figure 1:**
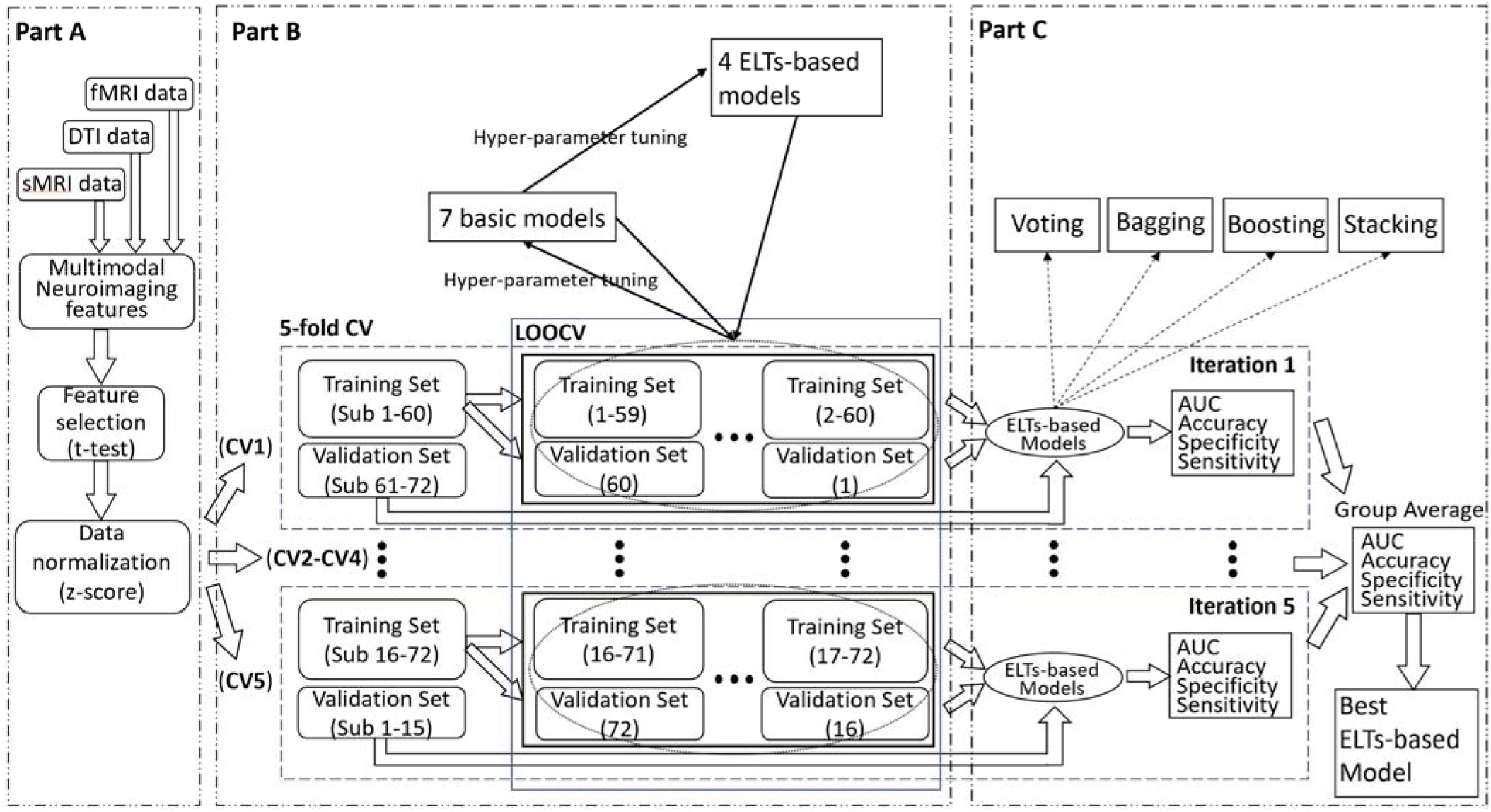
The ensemble learning flow chart. (sMRI: structural MRI; DTI: diffusion tensor imaging; fMRI: functional MRI; CV: cross-validation; LOOCV: leave-one-out cross-validation; AUC: the area under the receiver operating characteristic curve; ELTs: ensemble learning techniques)

We also applied unsupervised learning (i.e. the hierarchical clustering) in our dataset. The hyperparameters, including the metric used to compute the linkage (affinity), the linkage criterion used to determine which distance between sets of observation (linkage) were also tuned by using grid search. Then the model with best classification performance (i.e. accuracy) was selected.

### Regression Models

Following the results of the classification procedures, we also constructed the regression models to identify the relationships between the neuroimaging features and the clinical inattentive and hyperactive/impulsive symptoms T-scores. Based on the ELTs-based classification results, the top three important neuroimaging features were selected based on the weight of each feature in the optimal discriminators between ADHD and normal controls, as well as between ADHD persisters and remitters. Then, we applied Ordinary Least Squares (OLS) (Hutcheson, 1999), Ridge regression (Hoerl and Kennard, 1970), least absolute shrinkage and selection operator (LASSO) regression (Santosa and Symes, 1986; Tibshirani, 1996), Elastic Net regression (Zou and Hastie, 2005) to construct the prediction models for inattentive and hyperactive/impulsive t-scores, respectively. The same nested CV utilized in previous steps were also conducted in regression model construction. The hyperparameters included the regularization strength (alpha), solver to use in the computational routines (solver) for Ridge regression, the constant that multiplies the L1 term (alpha) for LASSO regression, the constant that multiplies the penalty terms (alpha), the Elastic Net mixing parameter (l1_ratio) for Elastic Net regression. During the iteration of 5-fold CV outer loop, the performance of each regression model with the optimal hyperparameters derived from LOOCV inner loop iterations was evaluated. The group average of regression performance, including the Pearson correlation coefficient and mean squared error (MSE) between predicted and observed values, of each regression model derived from each iteration of 5-fold CV were calculated.

### Evaluation Measures

The performance of each classification procedure classifier was measured in terms of classification accuracy, sensitivity, and specificity. The accuracy of a machine learning classification algorithm is to measure how often the algorithm classifies a data point correctly. It is defined as:

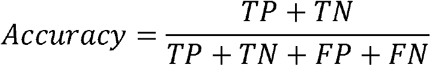

where *TP* is true positive, *TN* is true negative, *FP* is false positive, and *FN* is false negative.

Sensitivity describes the proportion of actual positive cases that are correctly identified as positive. It implies that there will be another proportion of actual positive cases, which would get predicted incorrectly as negative. The sensitivity is defined as:

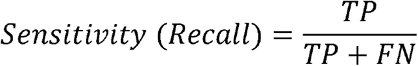

Specificity is a measure of the proportion of actual negatives, which got predicted as the negative. It implies that there will be another proportion of actual negative, which got predicted as positive and could be termed as false positives. It is defined as:

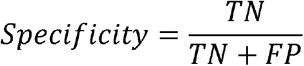

In addition, a ROC curve was plotted to illustrate the diagnostic ability of a binary classifier system as its discrimination threshold is varied. In the classification case, we calculated the confusion matrix for each iteration cycle of the classifier and calculated the AUC of ROC. AUC provides an aggregate measure of performance across all possible classification thresholds. One way of interpreting AUC is as the probability that the model ranks a random positive example more highly than a random negative example. The AUC of ROC is defined as:

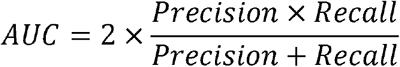

Among the equation of AUC, Precision and Recall are defined as:

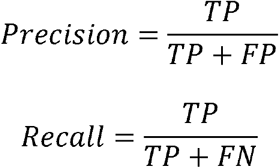

respectively.

For the regression model, the Pearson correlation coefficient and MSE between predicted values and actual values were calculated. The Pearson correlation coefficient is a measure of the linear correlation between two variables. It is defined as:

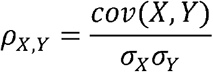

where *cov* is the covariance, σ_*X*_ is the standard deviation of *X*, σ_*y*_ is the standard deviation of *Y*.

The MSE of an estimator measures the average squared difference between the estimated values and the actual value, which is defined as:

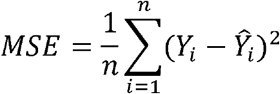

Where *X*_*i*_ and 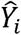 represent the actual and predicted value.

## Results

The demographic, clinical information and behavioral performance were summarized in Table 1. There were no significant demographic between-group differences. Moreover, all participants achieved a >85% rate for response accuracy when performing the fMRI task. Task performance measures, including reaction time, response accuracy rate, omission error rate, commission error rate did not show between-group differences (*p*>0.05).

The Table 3 **(Part I)** summarized the ADHD probands vs. controls classification performances of the basic models and ELTs. The classifier of SVM performed the best among all of those seven basic models regarding the AUC, accuracy, and specificity (AUC=0.87, accuracy=0.816, specificity=0.942). Furthermore, the bagging-based ELT with SVM as the basic model performed the best among all ensemble models (AUC = 0.89, accuracy=0.766, sensitivity=0.798). Meanwhile, the Table 3 **(Part II)** summarized the ADHD persisters vs. remitters classification performances of the basic models and ELTs, and again demonstrated that SVM performed the best among all the basic models regarding the AUC and accuracy (AUC=0.85, accuracy=0.7), while the bagging-based ELT with SVM as the basic model performed the best among all ensemble models (AUC = 0.9).

**Table 3:** The results of 7 basic and 4 ELTs-based classifications between the groups of ADHD and normal controls (**Part I**) as well as between the groups of ADHD persisters and ADHD remitters (**Part II**).

| Classifiers | Specificity | Sensitivity | Accuracy | AUC |
| --- | --- | --- | --- | --- |
| **Part I: ADHD vs. NC** |  |  |  |  |
| KNN | 0.72 | 0.66 | 0.689 | 0.69 |
| SVM | 0.942 | 0.69 | 0.816 | 0.87 |
| LR | 0.756 | 0.742 | 0.75 | 0.85 |
| NB | 0.778 | 0.718 | 0.748 | 0.86 |
| RF | 0.866 | 0.75 | 0.705 | 0.82 |
| LDA | 0.734 | 0.774 | 0.754 | 0.78 |
| MLP | 0.782 | 0.746 | 0.764 | 0.84 |
| ELT-Voting | 0.808 | 0.718 | 0.763 | 0.87 |
| ELT-Bagging | 0.734 | 0.798 | 0.766 | 0.89 |
| ELT-Boosting | 0.67 | 0.77 | 0.721 | 0.88 |
| ELT-Stacking | 0.756 | 0.742 | 0.75 | 0.82 |
| **Part II: ADHD-P vs. ADHD-R** |  |  |  |  |
| KNN | 0.4 | 0.934 | 0.667 | 0.72 |
| SVM | 0.65 | 0.75 | 0.7 | 0.85 |
| LR | 0.6 | 0.682 | 0.642 | 0.85 |
| NB | 0.734 | 0.65 | 0.692 | 0.77 |
| RF | 0.734 | 0.6 | 0.667 | 0.76 |
| LDA | 0.568 | 0.518 | 0.542 | 0.63 |
| MLP | 0.634 | 0.75 | 0.692 | 0.84 |
| ELT-Voting | 0.8 | 0.65 | 0.725 | 0.82 |
| ELT-Bagging | 0.75 | 0.582 | 0.67 | 0.90 |
| ELT-Boosting | 0.75 | 0.682 | 0.717 | 0.86 |
| ELT-Stacking | 0.884 | 0.684 | 0.783 | 0.78 |

ELT: ensemble learning technique; KNN: k-nearest neighbors; SVM: support vector machine; LR: logistic regression; RF: random forest; LDA: linear discriminant analysis; MLP: multilayer perceptron; AUC: the area under the receiver operating characteristic curve; ADHD: attention deficit/hyperactivity disorder; NC: normal controls; ADHD-R: ADHD remitters; ADHD-P: ADHD persisters.

The ROC curve for each classification procedure, including the unsupervised hierarchical clustering, in Figure 2., was also plotted. Results showed that classification performance parameters of the ELTs-based procedures were greatly improved compared to those of the basic model-based procedures. In addition, relative to the performance improvement between ensemble learning and basic models of the classification between ADHD and normal controls, the performance improvement of classification between ADHD persisters and remitter is much greater.

**Figure 2:**
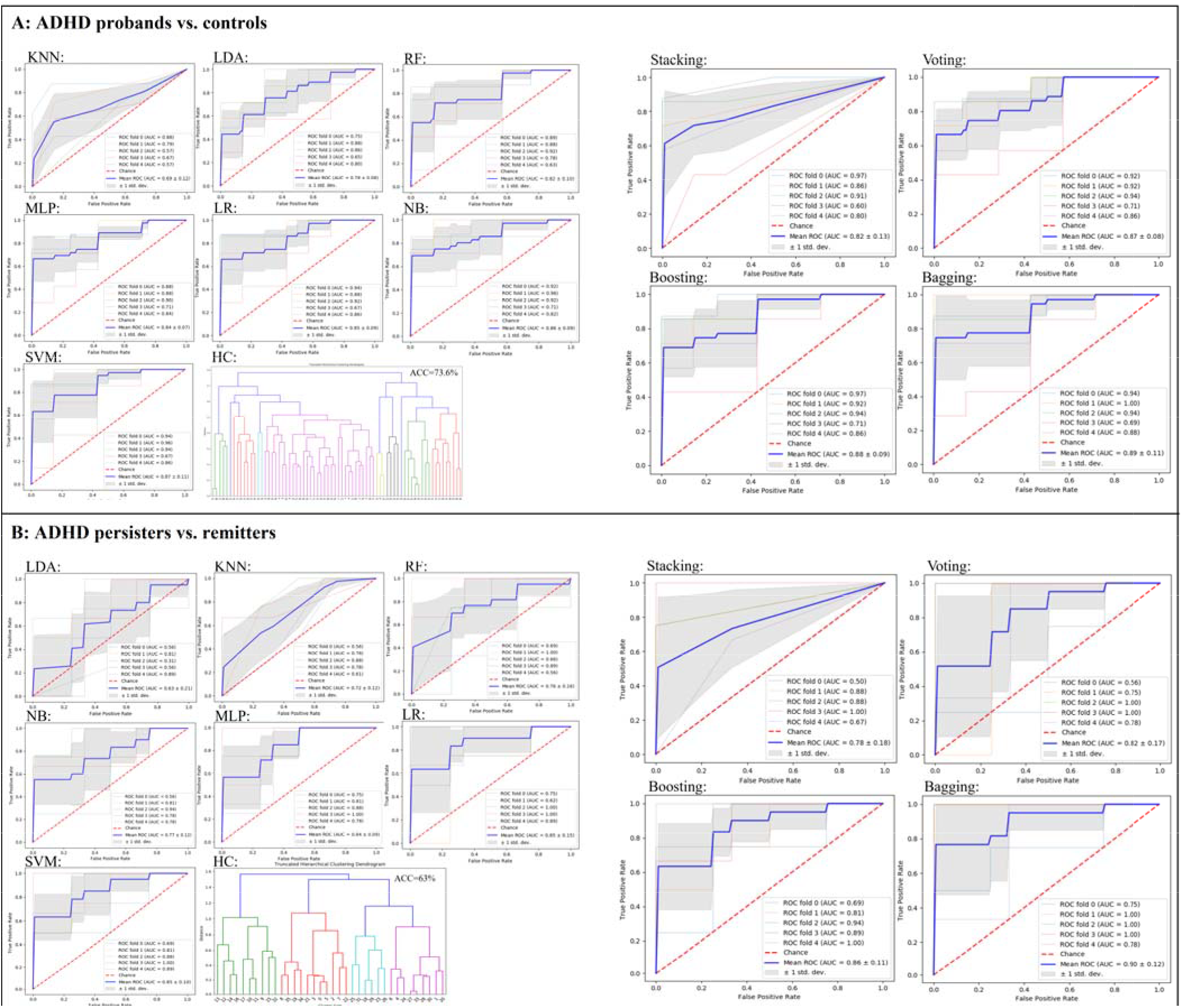
The AUC of each classification procedure for discrimination between ADHD probands and normal controls (**A**), and between ADHD persisters and remitters (**B**). (KNN: k-nearest neighbors; SVM: support vector machine; LR: logistic regression; RF: random forest; LDA: linear discriminant analysis; MLP: multilayer perceptron; HC: hierarchical clustering; ROC: the receiver operating characteristic curve; AUC: the area under the ROC curve)

The importance score of top three features for the classifications between ADHD probands and normal controls, and between ADHD persisters and remitters were showed in Table 4. More specifically, the nodal efficiency of right inferior frontal gyrus (IFG), the functional connectivity between right middle frontal gyrus (MFG) and right inferior parietal lobule (IPL), the volume of right amygdala served as the top three important features in the classification model between ADHD and normal controls. The nodal efficiency of right MFG, functional connectivity between right MFG and right IPL, and betweenness-centrality of left putamen played the top three important characteristics in the classification between ADHD persisters and remitters.

**Table 4:** The importance scores of top three features of classifications between ADHD probands and normal controls, as well as between ADHD persisters and ADHD remitters.

| Feature | Importance Score |
| --- | --- |
| **ADHD vs. NC** |  |
| Nodal efficiency of right Inferior Frontal gyrus | 0.134 |
| FC between right Middle Frontal gyrus and right Inferior Parietal lobule | 0.111 |
| Volume of right amygdala | 0.1 |
| **ADHD-P vs. ADHD-R** |  |
| Nodal efficiency of right Middle Frontal gyrus | 1.028 |
| FC between right Middle Frontal gyrus and right Inferior Parietal lobule | 0.852 |
| Betweenness-centrality of left putamen | 0.677 |

FC: functional connectivity; NC: normal controls; ADHD: attention deficit/hyperactivity disorder; ADHD-P: ADHD persists; ADHD-R: ADHD remitters.

The regression results (Table 5) showed that Elastic Net regression performed the best for the prediction of both inattentive and hyperactive/impulsive T-scores. Table 6 showed the importance scores of the top three features of Elastic Net regression for inattentive and hyperactive/impulsive symptom T-scores. In specific, the top three features for the prediction of inattentive T-score were the nodal efficiency of right IFG, the functional connectivity between MFG and IPL in right hemisphere, the volume of right amygdala. The top three features for the prediction of hyperactive/impulsive T-score included the nodal efficiency in right IFG, the functional connectivity between right MFG and right IPL, the nodal efficiency of right MFG.

**Table 5:** Pearson correlation coefficient and mean squared error performance of regression models.

| Regression | Pearson Correlation Coefficient | MSE |
| --- | --- | --- |
| **T-Inattentive** |  |  |
| OLS | $r=0.4603; p<0.001$ | 126.3 |
| LASSO | $r=0.4592; p<0.001$ | 124.6 |
| Ridge | $r=0.4605; p<0.001$ | 126.1 |
| Elastic Net | $r=0.4689; p<0.001$ | 121.1 |
| **T-Hyperactive/Impulsive** |  |  |
| OLS | $r=0.3329; p=0.0043$ | 126.5 |
| LASSO | $r=0.3395; p=0.0035$ | 123.3 |
| Ridge | $r=0.3334; p=0.0042$ | 126.3 |
| Elastic Net | $r=0.3488; p=0.0027$ | 119.8 |

OLS: ordinary least square; LASSO: least absolute shrinkage and selection operator; T-Inattentive: inattentive T-score; T-Hyperactive/Impulsive: Hyperactive/Impulsive T-score; MSE: mean squared error.

**Table 6:** The importance scores of top three features of Elastic Net regression for inattentive and hyperactive/impulsive symptom T-scores.

| Feature | r | p | Importance Score |
| --- | --- | --- | --- |
| **Inattentive** |  |  |  |
| Nodal efficiency of right Inferior Frontal gyrus | -0.399 | 0.001 | 3.471 |
| FC between right Middle Frontal gyrus and right Inferior Parietal lobule | 0.405 | <0.001 | 2.126 |
| Volume of right Amygdala | -0.011 | 0.928 | 1.819 |
| **Hyperactive/Impulsive** |  |  |  |
| Nodal efficiency of right Inferior Frontal gyrus | -0.345 | 0.003 | 2.289 |
| FC between right Middle Frontal gyrus and right Inferior Parietal lobule | 0.361 | 0.002 | 2.134 |
| Nodal efficiency of right Middle Frontal gyrus | -0.333 | 0.004 | 1.997 |

FC: functional connectivity.

## Discussion

To the best of our knowledge, this is the first study to apply ELTs in multimodal neuroimaging features generated from structural MRI, DTI, and task-based fMRI data collected from a sample of adults with childhood ADHD and controls, who have been clinically followed up since childhood. We found that the nodal efficiency in right IFG, functional connectivity between MFG and IPL in right hemisphere, and right amygdala volume were the most important features for discrimination between the ADHD probands and controls, while the nodal efficiency of right MFG, functional connectivity between right MFG and right IPL, and betweenness-centrality of left putamen played the most important roles for discrimination between the ADHD persisters and remitters. Moreover, the classification performance parameters of ELTs-based procedures were improved as compared to those of the basic classifiers.

Our current study observed the important roles of nodal efficiency in right IFG and functional connectivity between right MFG and right IPL for discrimination between ADHD probands and normal controls. The abnormalities of these regions were supported by a variety of existing neuroimaging and machine learning studies. Specifically, a substantial number of task-based and resting-state fMRI studies have consistently reported the decreased functional activation in right IFG (Cao et al., 2006; Konrad et al., 2006; Rubia et al., 2019; Rubia et al., 1999; Silk et al., 2005; Smith et al., 2006) and reduced functional connectivity between right MFG and right IPL (Lin et al., 2015; Vance et al., 2007) in children with ADHD as compared with normal controls. In addition, a substantial existing multivariate machine learning and ELTs-based studies have commonly reported that the functional activation and connectivity in frontal and parietal areas are associated with the classification performance between children with ADHD and normal controls (Brown et al., 2012; Colby et al., 2012; Deshpande et al., 2015; dos Santos Siqueira et al., 2014; Fair et al., 2012; Iannaccone et al., 2015; Qureshi et al., 2017). They have supported that the functional abnormalities in the frontal and parietal areas, which are critical components of the attention network in human brain, especially stimulus-driven top-down control, are associated with the symptom emergence of childhood ADHD (Posner and Rothbart, 2009). Moreover, we further found that the volume of right amygdala played a vital role in discrimination of ADHD probands and controls. The findings of amygdala anatomical abnormities in children with ADHD were supported by many previous studies. Amygdala played as a critically important role in emotion regulation (Banks et al., 2007; Davidson et al., 2000; Domes et al., 2010) and thus structural anomalies associated amygdala have been widely observed in children (Van Dessel et al., 2019) and adults with ADHD (Tajima-Pozo et al., 2018), which suggested that the aberrant structure of amygdala may be associated with less control of impulsivity and delay aversion (Van Dessel et al., 2018).

Additionally, our current study also indicated the important roles of nodal efficiency in right MFG, functional connectivity between right MFG and right IPL for discrimination between ADHD persisters and remitters and it was supported by a variety of existing neuroimaging studies. More specifically, reduced activations and functional connectivities in IFG, MFG, and fronto-parietal regions were observed in ADHD persisters when compared to ADHD remitters (Mattfeld et al., 2014; Schulz et al., 2017). The functional activations and connectivities in frontal and parietal regions during cognitive control were observed associated with the diverse adult outcomes of ADHD diagnosed in childhood, with symptom persistence linked to reduced activation or symptom recovery associated with higher connectivity within frontal areas (Francx et al., 2015; Mattfeld et al., 2014; Schulz et al., 2017).

Although several existing multivariate machine learning and ELTs-based studies have commonly reported that the anatomical features in frontal and parietal areas are associated with the classification performance between adults with ADHD and group-matched normal controls (Chaim-Avancini et al., 2017; Zhang-James et al., 2019), no machine learning study has been conducted to identify the classification pattern for discrimination between ADHD persisters and remitters. We further observed that the features of nodal efficiency in right IFG, functional connectivity between right MFG and right IPL, and right amygdala volume were associated with inattentive symptom severity T-score, while the nodal efficiencies of right IFG and MFG and functional connectivity between MFG and IPL in right hemisphere were associated with hyperactive/impulsive symptom severity T-score. These findings suggest the significant involvement of frontal and parietal lobes in right hemisphere for both inattentive and hyperactive symptom persistence of childhood ADHD (Francx et al., 2015).

Moreover, we also found that the classification performance parameters of ELTs-based procedures were improved compared to those of basic models. The ELTs have been developed in the big data science field to adaptively combine multiple basic classifiers in order to strategically deal with feature variance and bias, and optimize prediction performances (Balakrishnan et al., 2012; Dror et al., 2011; Hansen and Salamon, 1990; Schapire, 1990b). According to our classification results, bagging, sampling with replacement, would help to reduce the chance overfitting complex models. In our study, bagging with the basic model of SVC was applied to train our model and proved as the best classifier for both discriminations. In addition, we used AUC statistic for model evaluation, instead on commonly used accuracy, which can be influenced by case-control imbalance in data sets (Fawcett, 2006; Hanley and McNeil, 1982). Our study showed a satisfied performance of AUC with 0.89 and 0.9 for the discrimination between groups of ADHD and normal controls, and between the groups of ADHD persisters and remitters, respectively. Therefore, our study suggested that, as for the study with relatively small sample size, the ELTs-based models performed better than basic models.

In addition, the Elastic Net-based regression model demonstrated the best performance parameters when investigating the relations between the neuroimaging features and clinical symptom measures in the ADHD probands. The reason Elastic Net regression had the best performance was that it compromises the LASSO penalty (L1) and the ridge penalty (L2) (Zou and Hastie, 2005). The LASSO (L1) penalty function performs variable selection and dimension reduction by shrinking coefficients (Tibshirani, 1996), while the ridge (L2) penalty function shrinks the coefficients of correlated variables toward their average (Hoerl and Kennard, 1970). Therefore, as for the study with relatively small sample size, the combined method obviously performed better than isolated ones, e.g. LASSO regression and ridge regression.

In summary, together with existing findings, results of this study suggest that structural and functional alterations in frontal, parietal, and amygdala areas and their functional interactions significantly contributed to accurate discrimination of ADHD probands from controls; while abnormal fronto-parietal functional communications in the right hemisphere played an important role in symptom persistence in adults with childhood ADHD. In addition, the classification performance parameters (accuracy, AUC of the ROC, etc.) of ELTs-based procedures were greatly improved compared to those of basic model-based procedures.

Although the ELTs improved the model classification performances, especially for the cases when the base models had weak classification results, the current study has some limitations. First, our cohort consisted of both male and female subjects, but many more males. It is still unclear whether the discrimination models of ADHD differ between males and females. The future work may focus on constructing the classification models for both males and females. Second, the samples size of this study is relatively small. Although our study provided a considerable robust algorithm to reduce the overfitting, the relative small sample size may still influence the model’s performance. Future work will need a much larger cohort to test the ELTs.

## Acknowledgements

Dr. Xiaobo Li designed the study. Yuyang Luo managed literature searches, analyzed the clinical and imaging data, and wrote the first draft of the manuscript. Drs. Li, Alvarez, and Halperin edited and revised the manuscript. All authors contributed to and have approved the final manuscript.

