## Supplemental Table 1 for "Multimodal Neuroimaging-based Prediction of Adult Outcomes in Childhood-onset ADHD using Ensemble Learning Techniques"

Supplementary Table 1: Existing machine learning studies in neuroimaging data from children and/or adults with ADHD and group-matched controls.

| **Author, Year** | **Models** | **Validation** | **Features** | **Feature Type** | **Predictors** | **Acc** | **AUC** |
| --- | --- | --- | --- | --- | --- | --- | --- |
| **Children with ADHD** | | | | | | | |
| 1. (Brown et al., 2012) | SVM | 10-fold CV | rs-fMRI | Voxel | FC of bilateral thalamus, bilateral TL, MFG, PCC, Cerebellum | 0.71 | N/A |
| 2. (Colby et al., 2012) | SVM | Hold-out | sMRI, rs-fMRI | Voxel | FC, SA, cortical curvature in FL, CT and FC of CG and TL | 0.55 | N/A |
| 3. (Dai et al, 2012) | MKL | 10-fold CV | sMRI, rs-fMRI | ROI | N/A | 0.68 | 0.71 |
| 4. (Deshpande et al., 2015) | FCCANN | LOOCV | rs-fMRI | ROI | FC of OFC and cerebellum | 0.90 | N/A |
| 5. (Du et al., 2016) | SVM | 10-fold CV | rs-fMRI | Network | HSIC score of Operculum, Insula, putamen, STG | 0.95 | 0.97 |
| 6. (Eloyan et al., 2012) | Voting | Hold-out | sMRI, rs-fMRI | Voxel | FC between DM and DL in MC | 0.78 | N/A |
| 7. (Fair et al., 2012) | SVM | LOOCV | rs-fMRI | ROI | FC in PFC, PL, Cerebellum | 0.83 | N/A |
| 8. (Ghiassian et al., 2016) | MHPC | Hold-out | sMRI, rs-fMRI | Voxel | Voxel intensity and functional activation in FL, Cerebellum | 0.70 | N/A |
| 9. (Hart et al., 2014) | GPC | LOOCV | tb-fMRI | Voxel | Functional activation in PFC, CG, BG, Thalamus, PL | 0.77 | 0.81 |
| 10. (Iannaccone et al., 2015) | SVM | LOOCV | tb-fMRI | Voxel | Functional activation in SFG, PCC, TL, Brainstem, Cerebellum | 0.78 | 0.82 |
| 11. (Johnston et al., 2014) | SVM | LOOCV | sMRI | Voxel | Volume of Brainstem | 0.93 | N/A |
| 12. (Peng et al., 2013) | ELM | LOOCV | sMRI | ROI | SA and FI of FL, SA and FI in TL, SA and volume in OL, FI of Insula | 0.90 | 0.88 |
| 13. (Qureshi et al., 2016) | H-ELM | 10-by-10 Nested CV | rs-fMRI | ROI | N/A | 0.71 | N/A |
| 14. (Qureshi et al., 2017) | ELM | Random CV | sMRI, rs-fMRI | ROI | CT and FC in SFG and MTG | 0.93 | N/A |
| 15. (Sen et al., 2018) | SVM | Hold-out | sMRI, rs-fMRI | Voxel | N/A | 0.67 | N/A |
| 16. (Zou et al., 2017) | CNN | Hold-out | sMRI, rs-fMRI | Voxel | N/A | 0.69 | N/A |
| 17. (Yasumura et al., 2017) | SVM | 3-fold CV | fNIRS | ROI | Oxygenated hemoglobin change in PFC | 0.86 | 0.898 |
| 18. (Zhu et al., 2008) | PC-FDA | LOOCV | rs-fMRI | Voxel | ReHo of PFC, ACC, Thalamus | 0.85 | N/A |
| 19. (Zhang-James et al., 2019) | ELTs | Hold-out | sMRI | ROI | ICV, SA of FL, volume of Caudate and Thalamus | 0.61 | 0.67 |
| 20. (Kuang et al., 2014) | DBN | Hold-out | rs-fMRI | Voxel | N/A | 0.45 | N/A |
| 21. (Chang et al., 2012) | SVM | 3-fold CV | sMRI | ROI | N/A | 0.70 | N/A |
| 22. (Lim et al., 2013) | GPC | LOOCV | sMRI | Voxel | Voxel intensity in FL, Premotor, TL, Brainstem | 0.79 | 0.83 |
| 23. (Cheng et al., 2012) | SVM | LOOCV | rs-fMRI | ROI, Voxel | FC in FL and Cerebellum | 0.76 | N/A |
| **Adults with ADHD** | | | | | | | |
| 24. (Tenev et al., 2014) | Voting | 10-fold CV | EEG | ROI | N/A | 0.82 | N/A |
| 19. (Zhang-James et al., 2019) | ELTs | Hold-Out | sMRI | ROI | ICV, SA of FL, volume of Caudate and Thalamus | 0.62 | 0.66 |
| 25. (Chaim-Avancini et al., 2017) | SVM | 10-fold CV | sMRI, DTI | ROI, Voxel | GM and WM intensity across FL, TL, OL, Thalamus, Cerebellum, FA | 0.66 | 0.71 |

ICV: intracranial volume; MFG: middle frontal gyrus; MC: motor cortex; SMC: sensorimotor cortex; PFC: prefrontal cortex; FL: frontal lobe; PL: parietal lobe; TL: temporal lobe; OL: occipital lobe; BG: basal ganglia; CG: cingulate gyrus; MTG: middle temporal gyrus; OFC: orbitofrontal cortex; STG: superior temporal gyrus; PFC: prefrontal cortex; MHPC: (f)MRI HOG-feature-based patient classification; GPC: Gaussian process classifiers; ACC: anterior cingulate cortex; PCC: posterior cingulate cortex; H-ELM: hierarchical extreme learning machine; ELM: extreme learning machine; DBN: deep belief network; CNN: convolutional neural network; MKL: multiple kernel learning; PC-FDA: principle component-based Fisher discriminative analysis; ROI: region of interest; Acc: accuracy; CV: cross validation; LOOCV: leave-one-out cross validation; sMRI: structural magnetic resonance imaging; rs-fMRI: resting-state functional magnetic resonance imaging; tb-fMRI: task-based functional magnetic resonance imaging; FC: functional connectivity; SA: surface area; CT: cortical thickness; FI: folding index; ReHo: regional homogeneity; HSIC: Hilbert-Schmidt Independence Criterion; DM: dorsomedial; DL: dorsolateral; GM: gray matter; WM: white matter; N/A: not available.
